# Nrf2 is a central regulator of the metabolic landscape in macrophages and finetunes their inflammatory response

**DOI:** 10.1101/2021.08.13.456204

**Authors:** Dylan G. Ryan, Elena V. Knatko, Alva Casey, Jens L. Hukelmann, Alejandro J. Brenes, Sharadha Dayalan Naidu, Maureen Higgins, Laura Tronci, Efterpi Nikitopoulou, Luke A.J. O’Neill, Christian Frezza, Angus I. Lamond, Andrey Y. Abramov, Doreen A. Cantrell, Michael P. Murphy, Albena T. Dinkova-Kostova

## Abstract

To overcome oxidative, inflammatory, and metabolic stress, cells have evolved networks of cytoprotective proteins controlled by nuclear factor erythroid 2 p45-related factor 2 (Nrf2) and its main negative regulator the Kelch-like ECH associated protein 1 (Keap1). Here, we used high-resolution mass-spectrometry to characterize the proteomes of macrophages with genetically altered Nrf2 status. Our analysis revealed significant differences among the genotypes in cellular metabolism and redox homeostasis, which we validated with respirometry and metabolomics, as well as in anti-viral immune pathways and the cell cycle. Nrf2 status significantly affected the proteome following lipopolysaccharide (LPS) stimulation, with alterations in redox, carbohydrate and lipid metabolism, and innate immunity observed. Of note, Nrf2 activation was found to promote mitochondrial fusion in inflammatory macrophages. The Keap1 inhibitor, 4-octyl itaconate (4-OI), a derivative of the mitochondrial immunometabolite itaconate, remodeled the inflammatory macrophage proteome, increasing redox and suppressing anti-viral immune effectors in a Nrf2-dependent manner. These data suggest that Nrf2 activation facilitates metabolic reprogramming and mitochondrial adaptation, and finetunes the innate immune response in macrophages.

**Graphical abstract:** 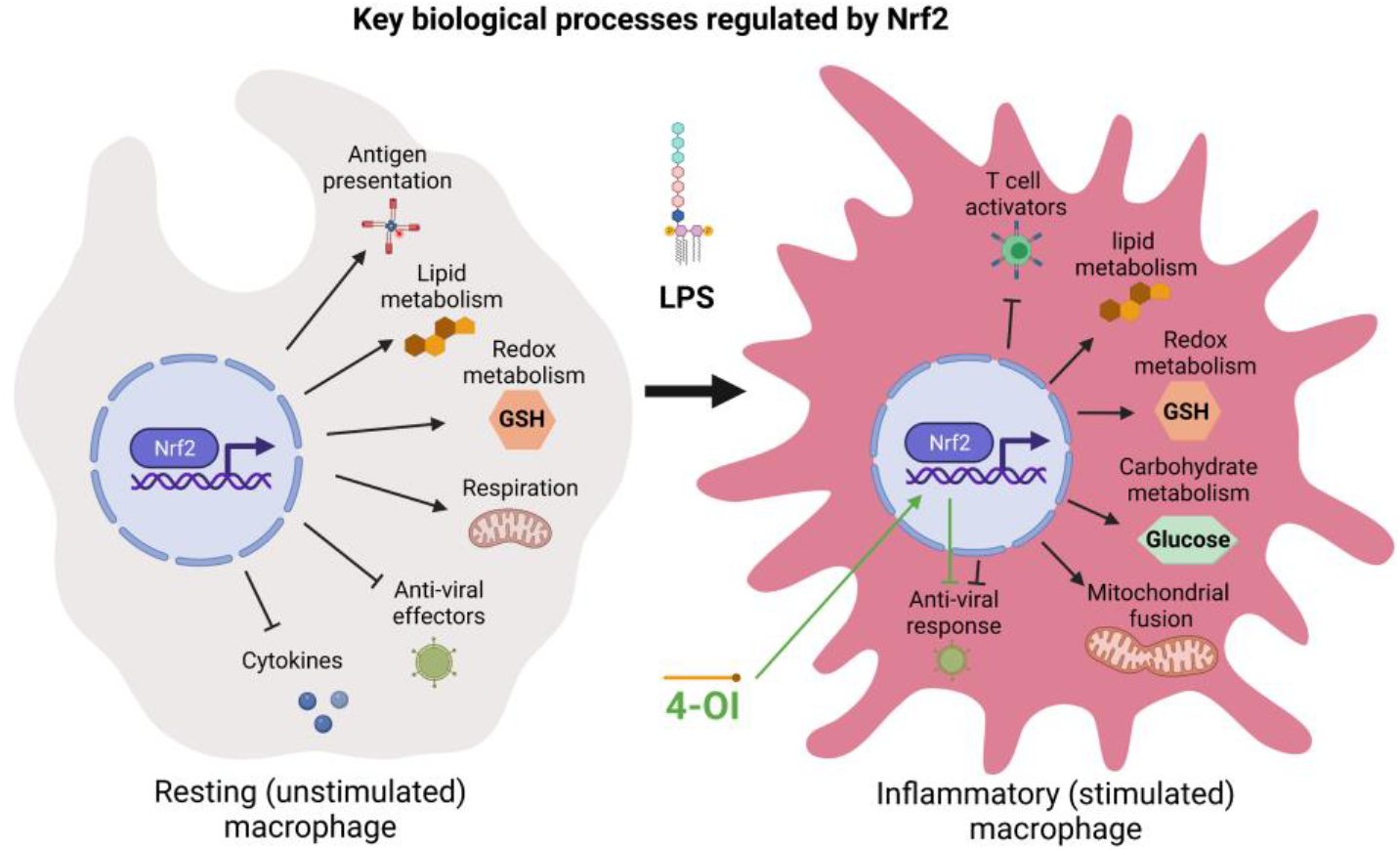

**Highlights:**

- First high-resolution proteome of macrophages with genetically altered Nrf2 status
- Nrf2 is key regulator of macrophage redox and intermediary metabolism
- Nrf2 finetunes the inflammatory response suppressing anti-viral immune and cytokine effectors, whilst promoting T cell activation factors
- Nrf2 regulates mitochondrial adaptation in inflammatory macrophages promoting the formation of a fused network
- 4-octyl itaconate (4-OI) suppresses anti-viral immune effectors in inflammatory macrophages in a Nrf2-dependent manner

## Introduction

The transcription factor nuclear factor erythroid 2-related factor 2 (Nrf2, gene name *Nfe2l2*), and its negative regulator Kelch-like ECH associated protein 1 (Keap1) are at the interface of redox and intermediate metabolism (Hayes and Dinkova-Kostova, 2014; Yamamoto et al., 2018), and have a complex, but incompletely understood, function in infection, inflammation and immunity (Cuadrado et al., 2020). This is not surprising considering that infection and inflammation cause disturbances in cellular redox homeostasis, which is restored by upregulation of Nrf2-target proteins (Hayes and Dinkova-Kostova, 2014). 4-octyl itaconate (4-OI), a derivative of the immunometabolite itaconate that activates Nrf2 via Keap1 alkylation, suppresses certain pro-inflammatory cytokines in macrophages in vitro and is protective in a LPS lethality model in vivo (Mills et al., 2018). Furthermore, genetic and pharmacological Nrf2 activation is considered anti-inflammatory and facilitates the resolution of inflammation (Dayalan Naidu et al., 2018; Kobayashi et al., 2016).

Nrf2 also activates transcription of genes important for macrophage function, such as macrophage receptor with collagenous structure (MARCO) (Harvey et al., 2011), a receptor required for bacterial phagocytosis, cluster of differentiation 36 (CD36) (Maruyama et al., 2008), a scavenger receptor for oxidized low-density lipoproteins, and the virus surveillance mediator interleukin-17D (IL-17D) (Saddawi-Konefka et al., 2016). In cancer cells, Nrf2 promotes replication of the vesicular stomatitis virus Δ51, facilitating oncolytic infection (Olagnier et al., 2017). By contrast, Nrf2 is inactivated by herpes simplex virus 1 (HSV-1) or severe acute respiratory syndrome coronavirus 2 (SARS-CoV-2), while the Nrf2 activators sulforaphane and 4-OI inhibit replication of these viruses, correlating with increased resistance to infection (Olagnier et al., 2020; Ordonez et al., 2021; Wyler et al., 2019).

Although the downregulation of pro-inflammatory responses by Nrf2 activation is consistently observed in cells and in animal and human tissues (Dayalan Naidu et al., 2018; Knatko et al., 2015; Kobayashi et al., 2016; Liu et al., 2020; Thimmulappa et al., 2006), global in-depth insights of the effect of Nrf2 on macrophages and their responses to inflammatory stimuli are lacking. In this study, we present the first analysis of how Nrf2 affects the proteome and metabolome of differentiated macrophages, comparing both resting and stimulated states. We identified a central role for Nrf2 in regulating the metabolic landscape of macrophages, influencing a plethora of processes involved in redox, carbohydrate and lipid metabolism. Moreover, we found a role for Nrf2 in regulating respiration and mitochondrial fusion in activated macrophages. Additionally, Nrf2 was found to have divergent effects on immune effectors, suppressing anti-viral response proteins but promoting factors involved antigen presentation. Finally, we identified a crucial role for Nrf2 in mediating the suppressive effects of 4-OI on interferon effector proteins and the activation of redox and intermediary metabolic enzymes. These findings place Nrf2 at the center of macrophage metabolism and has major implications for our understanding of Nrf2 biology and how it regulates the innate immune response.

## Results

### The proteomes of resting and activated macrophages with altered Nrf2 status

To understand how Nrf2 regulates macrophage biology, we used high-resolution mass-spectrometry (MS) to characterize the proteomes of differentiated bone marrow-derived macrophages (BMDMs) isolated from, respectively, wild-type (WT), Keap1-knockdown (Keap1-KD, expressing ~70% lower levels of Keap1 compared to WT, and consequently high Nrf2 levels) and Nrf2-knockout (Nrf2-KO, expressing transcriptionally inactive disrupted Nrf2-LacZ fusion protein) mice (Knatko et al., 2020) both in a resting (unstimulated) state and following LPS stimulation (100 ng/ml, 24 h) **(Figure 1A**). We confirmed the genotype of the macrophages by immunoblotting for Nrf2, Keap1, and the prototypical Nrf2 target, Nqo1 (**Figure 1B**). Interestingly, we observed some small but significant changes in cell volume among the genotypes, whereby Nrf2-KO and Keap1-KD resulted in a decrease and an increase, respectively (**Figure S1A**).

**Figure 1.**
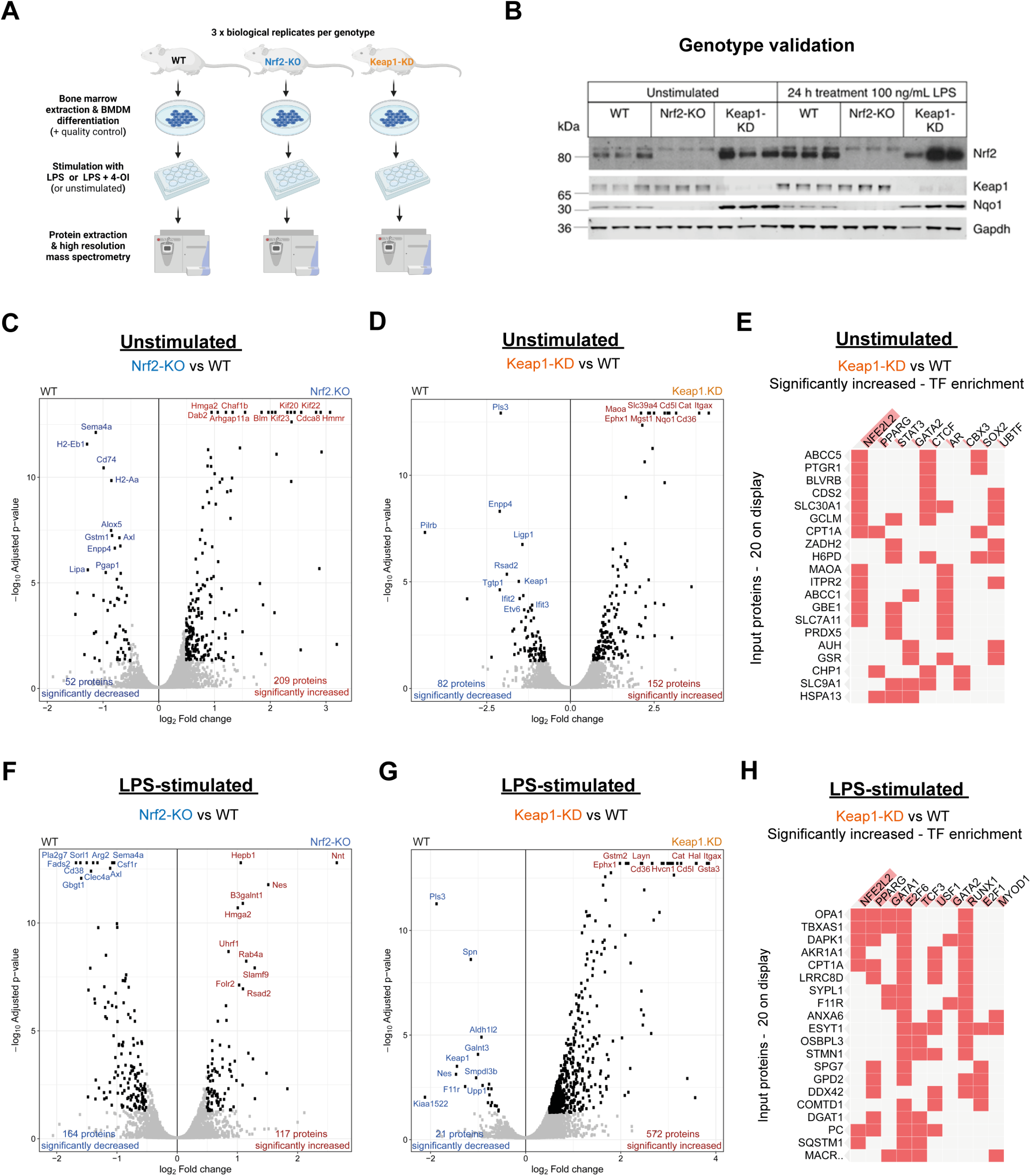
Nrf2 remodels the proteome in resting and activated macrophages. **(A)** Experimental design and workflow **(B)** Validation of WT, Nrf2 and Keap1-KO genotypes (n = 3 biological replicates). **(C)** Volcano plot of unstimulated Nrf2-KO compared to WT **(D)** Volcano plot of unstimulated Keap1-KD compared to WT **(E)** Transcription factor (TF) enrichment of increased targets in unstimulated Keap1-KD compared to WT **(F)** Volcano plot of LPS-stimulated Nrf2-KO compared to WT **(G)** Volcano plot of LPS-stimulated Keap1-KD compared to WT **(H)** TF enrichment of increased targets in LPS-stimulated Keap1-KD compared to WT **(C-D, F-G)** (n = 3 biological replicates). Cut-offs - log_2_FC = 0.5; FDR < 0.05, determined using t statistics. **(E, F)** Determined by Enrichr using ENCODE and ChEA database (combined score – p value and z score).

The levels of Nrf2 increased following macrophage activation (**Figure 1B**), consistent with previous findings (Mills et al., 2018). Differential expression analysis in the Nrf2-KO and Keap1-KD genotypes revealed substantial changes to the proteome in both resting and activated macrophages (**Figure 1C-D & 1F-G**). Nrf2 targets were enriched in the significantly increased proteins of Keap1-KD macrophages (**Figure 1E, H and Figure S1B**). Similarly, transcription factor (TF) enrichment analysis of proteins significantly decreased with Nrf2-KO confirmed enrichment for Nrf2 targets (**Figure S1C**). Of note, Nrf2 disruption led to a greater decrease in the number of proteins in LPS-stimulated macrophages (**Figure 1F**), whereas Keap1-KD led to a substantial increase (**Figure 1G**), which suggests that Nrf2 stabilization is an important feature of the LPS response. Enrichment for innate immune pathways, including cytokine and interferon signaling, was detected in LPS-activated WT macrophages (**Figure S1D**), and LPS increased the transcript levels of *Il1b* (**Figure S1E**), validating the ability of LPS to promote macrophage activation. Furthermore, we confirmed the ability of 4-OI to inhibit *Il1b* in part via Nrf2 (**Figure S1E**). Finally, Keap1-KD led to a suppression of *Il1b*, showing that genetic activation of Nrf2 also suppresses specific pro-inflammatory cytokines.

### Nrf2 is critical regulator of metabolism and innate immune pathways in resting macrophages

To better understand the biological processes that Nrf2 regulates in the resting state, we performed an over-representation analysis (ORA) on the significantly decreased (Nrf2 positive regulation) (**Figure 2A and Figure S2A**) and increased (Nrf2 negative regulation) (**Figure S2D**) proteins of Nrf2-KO macrophages. The analysis of positively regulated processes revealed a large functional cluster, which included antigen processing and presentation and factors involved in regulating T cell responses (**Figure 2A**). Lipid and glutathione (GSH) metabolism, as well as positive regulators of type 2 immune response, were also found to increase (**Figure S2A**). Amongst the significantly increased proteins, two functional clusters were observed, which included proteins involved in cell division and DNA replication and repair (**Figure S2D**).

**Figure 2.**
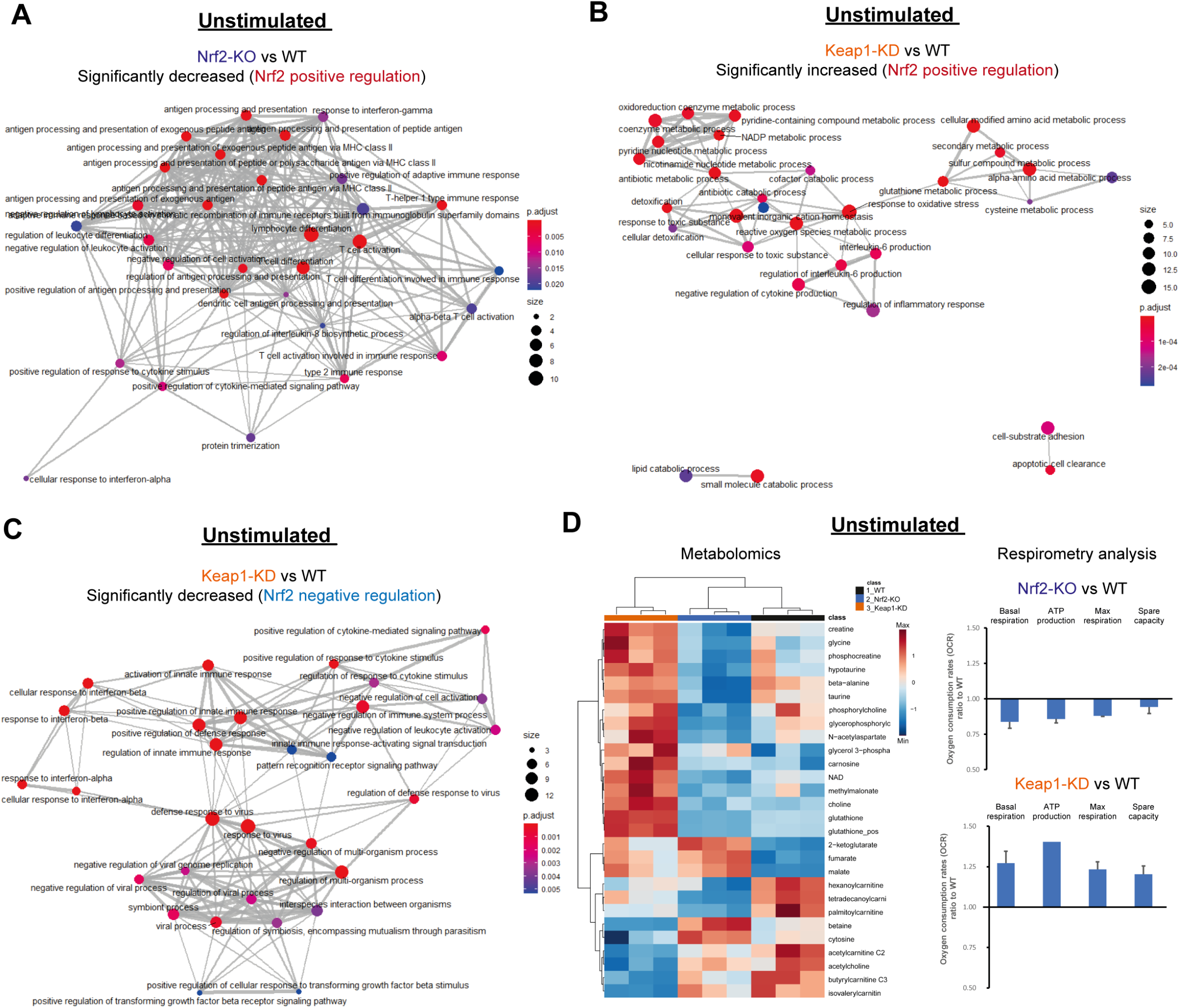
Nrf2 suppresses proteins involved in anti-viral immunity and cytokine production, while maintaining cellular redox metabolism. **(A)** Enrichment map of GO: biological processes of Nrf2 positively regulated targets (Nrf2-KO vs WT) **(B)** Enrichment map of GO: biological processes of Nrf2 positively regulated targets (Keap1-KD vs WT) **(C)** Enrichment map of GO: biological processes of Nrf2 negatively regulated targets – Keap1-KD versus WT **(A-C)** ORA by clusterProfiler, FDR correction by Bonferroni test. **(D)** Heatmap of significantly altered metabolites (n = 3 biological replicates) and oxygen consumption rates (OCR) (representative of 3 biological replicates). p value determined by one-way ANOVA, corrected for multiple comparisons by Tukey statistical test. Cut off – FDR < 0.05.

In Keap1-KD macrophages, ORA of the significantly increased proteins (Nrf2 positive regulation) identified three functional clusters (**Figure 2B**). The major cluster included biological processes involved in oxidative stress, sulfur metabolism and inflammatory response regulators, while the final two clusters identified were lipid catabolism and cell adhesion (**Figure 2B**). The most significantly enriched pathways included metabolism, biological oxidations and GSH-mediated detoxification (**Figure S2B**). In contrast, ORA of negatively regulated processes revealed two interconnected functional clusters, including proteins involved in anti-viral immunity and positive regulators of the innate immune response (**Figure 2C**). Notably, interferon signaling was the most enriched downregulated pathway with Nrf2 activation (**Figure S2C**). Thus, our proteomic analysis suggests that Nrf2 is required to maintain cellular redox and lipid homeostasis, while modulating distinct innate immune effector functions.

To validate a role for Nrf2 in regulating metabolism, we performed liquid chromatography-mass spectrometry (LC-MS)-based metabolomic analysis of resting WT, Nrf2-KO and Keap1-KD macrophages (**Figure 2D, left panel**). A clear segregation of the metabolome according to genotype was observed (**Figure 2D and Figure S2E**), validating a role for Nrf2 in regulating basal macrophage metabolism. Notably, Nrf2 activation significantly increased antioxidant metabolites, including GSH, hypotaurine, taurine, β-alanine and carnosine, whereas Nrf2 disruption significantly decreased intracellular GSH, taurine, hypotaurine and β-alanine (**Figure 2C**). Indeed, the rate-limiting GSH biosynthetic enzymes, Gclc and Gclm, as well as Nqo1 and Gsr, were decreased with Nrf2-KO and increased with Keap1-KD (**Figure S2F**). Furthermore, both Nrf2 activation and disruption led to significant alterations in mitochondrial metabolites, such as those involved in fatty acid oxidation (FAO) (carnitine, palmitoylcarnitine, hexanoylcarnitine and tetradecanoylcarnitine), the TCA cycle (fumarate, malate and 2-ketoglutarate) and bioenergetics (NAD, creatine, phosphocreatine). These findings suggest an involvement of Nrf2 in regulating mitochondrial metabolism in macrophages.

To confirm these observations using MS-independent methods, we performed respirometry analysis of oxygen consumption rates (OCR) in all three genotypes. This identified a role for Nrf2 in regulating mitochondrial respiration (**Figure 2D, right panel**). Nrf2 activation increased the basal respiration rates associated with ATP production, in agreement with the changes observed in the metabolome and previous experiments in mouse embryonic fibroblasts, neurons, and isolated mitochondria (Holmstrom et al., 2013; Ludtmann et al., 2014). On the other hand, Nrf2 disruption decreased respiration, and the above respiration-associated parameters (**Figure 2D, right panel**).

In summary, these results support an essential role for Nrf2 in governing redox and intermediary metabolism in resting macrophages, and other cellular processes, such as the innate immune response and DNA damage repair.

### Nrf2 is critical regulator of metabolism, mitochondrial adaptation and innate immune pathways in inflammatory macrophages

To better understand the biological processes that Nrf2 regulates during an inflamed state, we performed an ORA on the significantly decreased (**Figure 3A**) and increased (**Figure S3A**) proteins in Nrf2-KO macrophages stimulated with LPS. Enrichment analysis of positively regulated processes identified two functional clusters i.e. a large cluster that includes a plethora of intermediary metabolic pathways associated with carbohydrate, cofactor and energy metabolism, and a smaller cluster for the cellular response to oxidative stress (**Figure 3A**). The most significantly enriched pathways included those involved in glycolysis and GSH metabolism (**Figure S3C**). In Keap1-KD positively regulated processes, a loosely interconnected functional cluster was observed with an enrichment in processes associated with lipid metabolism, amino acid metabolism and cofactor metabolism, while enrichment in regulators of reactive oxygen species (ROS), cell adhesion and organic ion transport were also observed (**Figure 3B**). The most significantly enriched pathways included those related to lipid metabolism, such as FAO, and ROS detoxification (**Figure S3D**). In contrast, we did not observe many significant increases in the proteome of LPS-treated Nrf2-KO macrophages with no clearly regulated pathways (**Figure S3A**), whilst a large functional cluster was associated with the regulation of T cell activation in the decreased targets of Keap1-KD macrophages (**Figure S3B**).

**Figure 3.**
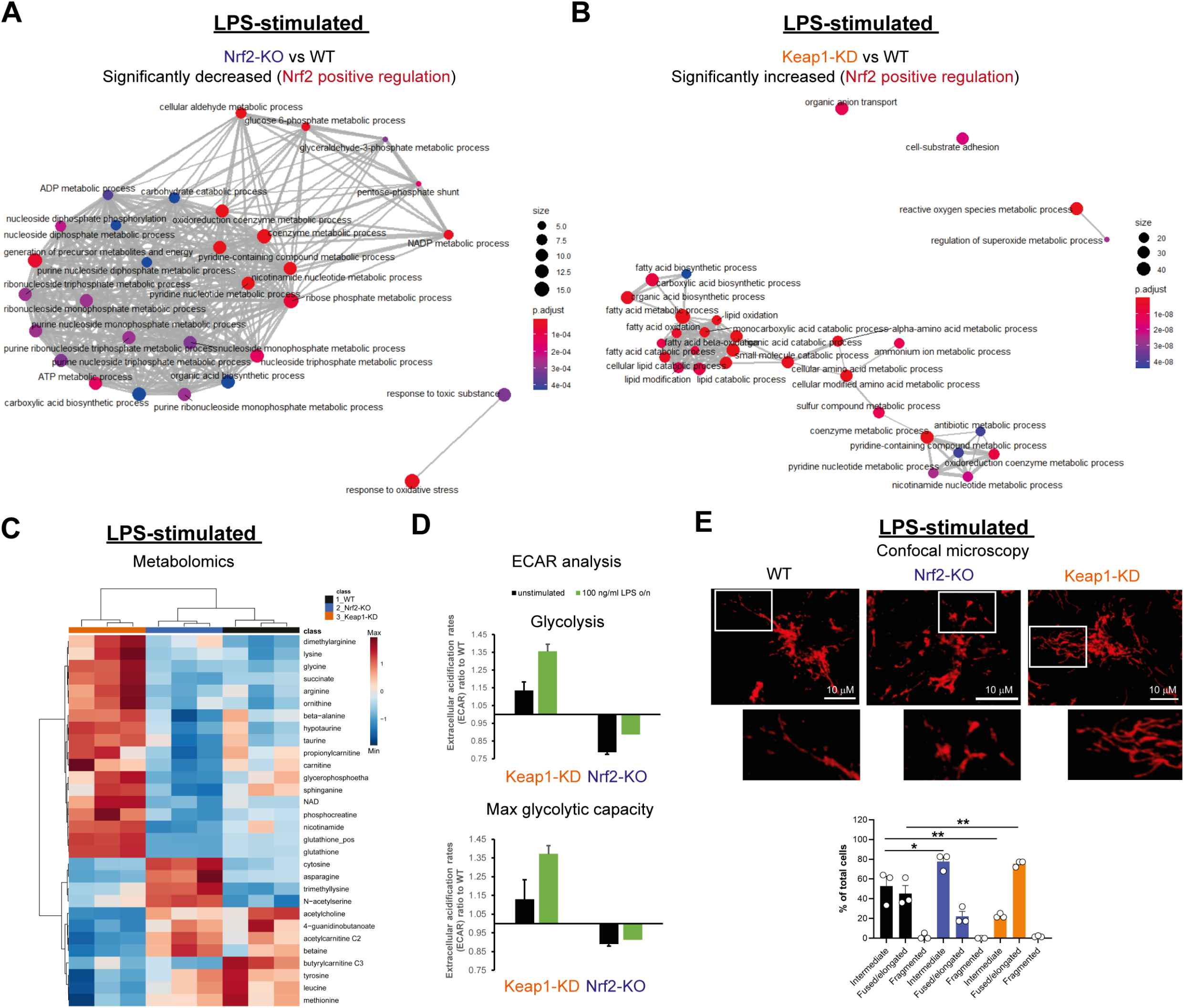
Nrf2 is a central regulator of metabolism, mitochondrial adaptation and immune effector functions in inflammatory macrophages. **(A)** Enrichment map of GO: biological processes of Nrf2 positively regulated targets (Nrf2-KO vs WT) **(B)** Enrichment map of GO: biological processes of Nrf2 positively regulated targets (Keap1-KD vs WT) **(A-B)** ORA by clusterProfiler, FDR correction by Bonferroni test. **(C)** Heatmap of significantly altered metabolites (n = 3 biological replicates). p value determined by one-way ANOVA, corrected for multiple comparisons by Tukey statistical test. Cut off – FDR < 0.05. **(D)** Extracellular acidification rates (ECAR) (representative of 3 biological replicates) **(E)** Confocal microscopy of mitochondrial morphology using TOM20 (images are representative, bar plot n = 3 biological replicates). Data are mean ± SEM. p value determined by one-way ANOVA, corrected for multiple comparisons by Tukey statistical test. p <0.05*; p <0.01**; p < 0.001***.

LC-MS-based metabolomic analysis of WT, Nrf2-KO and Keap1-KD macrophages stimulated with LPS (**Figure 3C**) revealed a significant increase in metabolites associated with the antioxidant response and bioenergetics. We also observed significant alterations in the abundance of several amino acids, with increased asparagine levels in Nrf2-KO, while tyrosine, leucine and methionine were decreased, and arginine, lysine and glycine were increased in the Keap1-KD cells. These findings support a central role for Nrf2 in governing macrophage metabolism, as predicted by our proteomic analyses (**Figure 3A-B & S3C-D**). Likewise, analysis of extracellular acidification rates (ECAR) also confirmed a role for Nrf2 in promoting glycolysis in resting and activated macrophages (**Figure 3D**).

Interestingly, in addition to changes in metabolic enzymes, we also observed a significant increase in the mitochondrial fusion proteins, Opa1, Mfn1 and Mfn2, with Keap1-KD and a significant increase in the mitochondrial fission factors, Mff and Mief2, with Nrf2-KO in activated macrophages (**Figure S3F**). Given the importance of mitochondrial physiology in governing cellular redox state, metabolism and bioenergetics, and the clear regulation of these processes by Nrf2, we hypothesized that Nrf2 status may modulate mitochondrial morphology. To explore this, we performed confocal microscopy analysis of mitochondrial morphology following immunofluorescence staining of the outer mitochondrial membrane (OMM) protein Tom20 (**Figure 3E and Figure S3E**). Mitochondrial morphology was assigned as intermediate, fused/elongated or fragmented (Tabara and Prudent, 2020). In the unstimulated state, mitochondria predominantly exhibited intermediate morphology across all three genotypes, however, a minority of Keap1-KD mitochondria exhibited fragmented or fused/elongated morphologies (**Figure S3E**). Interestingly, 24 h of LPS stimulation in WT macrophages caused a notable change, with 45% of cells displaying fused/elongated morphology, while the percentage of cells with intermediate morphology decreased from 95- to 55% (**Figure 3E and Figure S3E**). The LPS-mediated change in mitochondrial morphology was even more striking in Keap1-KD cells, where the percentage of cells with intermediate mitochondria decreased from 75- to 25%, whereas the percentage of cells with fused/elongated morphology increased, from 5- to 75% (**Figure 3E and Figure S3E**). In contrast, LPS treatment of Nrf2-KO cells resulted in only a modest increase in fused/elongated mitochondria (10 to 25%), whereas the percentage of cells with intermediate mitochondria decreased from 90- to 75% (**Figure 3E and Figure S3E**). Together, these experiments illustrate that prolonged stimulation of macrophages with LPS causes a switch in mitochondrial morphology, from intermediate to fused/elongated, which is enhanced by Nrf2 activation and suppressed by Nrf2 disruption. Therefore, Nrf2 represents a crucial factor governing redox and intermediary metabolism, mitochondrial adaptation and innate immunity in macrophages upon encountering infectious stimuli.

### The Keap1 inhibitor, 4-octyl itaconate, promotes redox metabolism and inhibits anti-viral immune effectors in inflammatory macrophages

During LPS stimulation, macrophages undergo profound metabolic changes, engaging aerobic glycolysis and suppressing OXPHOS (Ryan et al., 2019; Ryan and O’Neill, 2020). Importantly, several mitochondrial metabolites, including succinate, fumarate and itaconate accumulate and act as signals to regulate macrophage effector functions (Mills et al., 2018; Ryan et al., 2019). Our previous work demonstrated that a lipophilic derivative of itaconate, 4-OI, is a robust Nrf2 activator and anti-inflammatory compound (Mills et al., 2018). 4-OI activates Nrf2 via alkylation of key cysteines on Keap1, and this is, at least in part, responsible for its anti-inflammatory effects (**Figure S1E)**. In the same study, we found that 4-OI inhibited Interferon-β production and the expression of interferon-inducible targets, but the role of Nrf2 was unclear. Therefore, we performed proteomic analysis of LPS-stimulated WT and Nrf2-KO macrophages that had been treated with 4-OI to determine to what extent Nrf2 was involved in mediating remodeling of the macrophage proteome upon treatment of 4-OI (**Figure S4A-C**). Indeed, we observed significant changes in the proteome of activated macrophages treated with 4-OI in both genotypes; however, the impact was far more pronounced in WT cells (**Figure S4A-B**). We also confirmed an enrichment for Nrf2 targets in WT cells treated with 4-OI (**Figure S4C**).

To understand what biological processes 4-OI regulates, we performed ORA on the significant changes, and found that in WT macrophages 4-OI regulates four functional clusters (**Figure 4A**). Unsurprisingly, an enrichment for redox metabolism, detoxification and lipid metabolism was identified, while an increase in positive regulators of cytokine production also emerged. 4-OI significantly decreased anti-viral immune effectors (**Figure 4B,E**) and positive regulators of leukocyte activation, such as Nos2 (**Figure S4A**), consistent with its reported anti-inflammatory role (Mills et al., 2018). Strikingly, in Nrf2-KO macrophages, there was no biological processes that reached significance and 4-OI lost its ability to regulate both redox metabolism and certain immune response effectors (**Figure 4C**). Of note, 4-OI lost its ability to suppress anti-viral immune effectors and other immune proteins in Nrf2-KO macrophages (**Figure 4D**). This highlights an important role for Nrf2 in mediating 4-OI-induced remodeling of the proteome and mediating aspects of its immunomodulatory capabilities.

**Figure 4.**
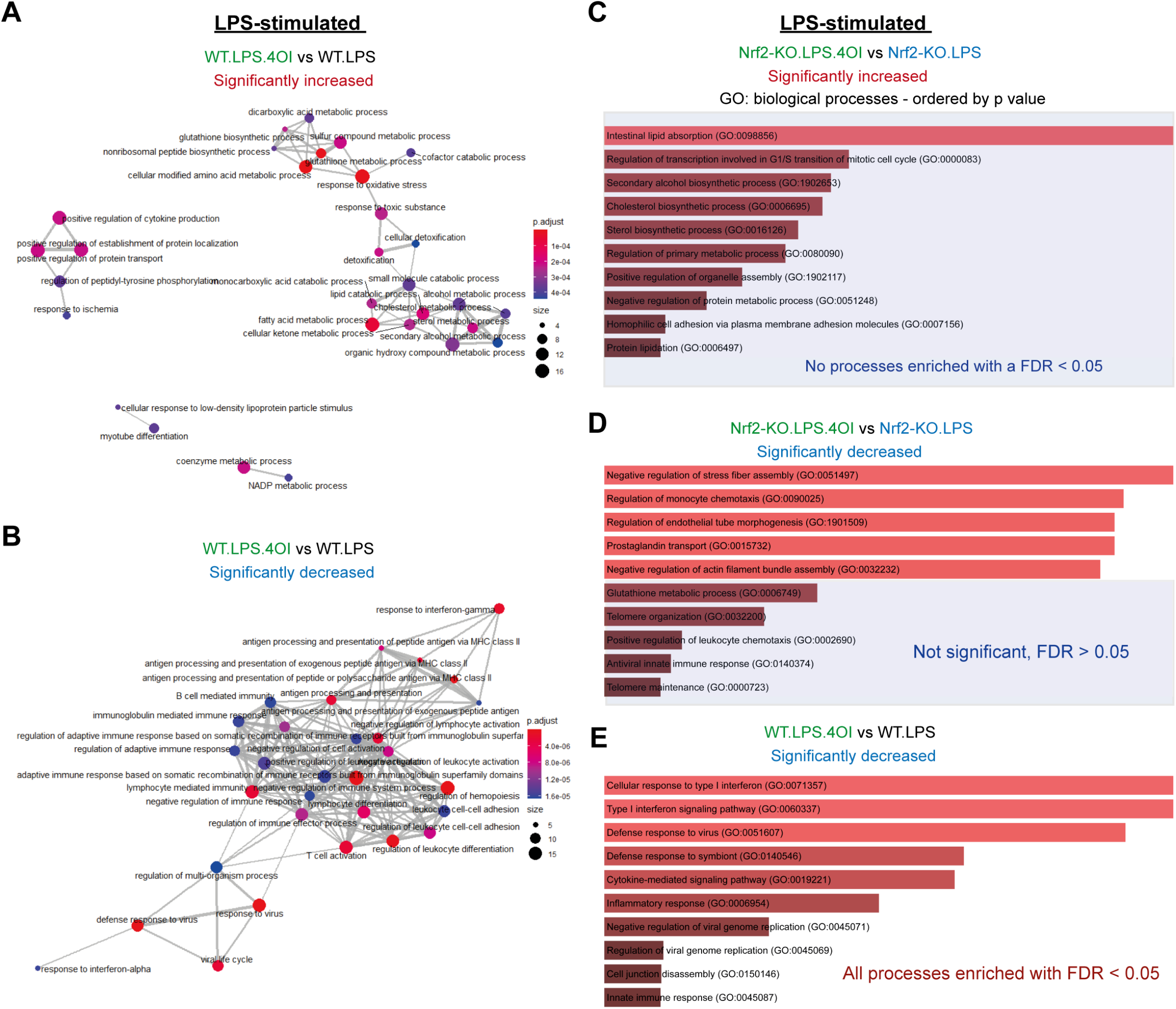
4-OI regulates redox metabolism and suppresses anti-viral responses in a Nrf2-dependent manner. **(A)** Enrichment map of GO: biological processes of 4-OI positively regulated targets (WT) **(B)** Enrichment map of GO: biological processes of 4-OI negatively regulated targets (WT) **(C)** Enrichment map of GO: biological processes of 4-OI positively regulated targets (Nrf2-KO) **(A-C)** ORA by clusterProfiler, FDR correction by Bonferroni test. **(D)** Enrichment of GO: biological processes of 4-OI negatively regulated targets (Nrf2-KO). **(E)** Enrichment of GO: biological processes of 4-OI negatively regulated targets (WT). **(A-E)** 4-OI used at 125 μM (**D-E**) ORA by Enrichr and FDR correction by Bonferroni test.

## Discussion

The cytoprotective Keap1-Nrf2 axis is the master regulator of the antioxidant stress response in mammals (Hayes and Dinkova-Kostova, 2014). Importantly, Nrf2 status has clinical implications for infectious and metabolic disease, and autoimmunity, which is only beginning to be understood. Nrf2 is stabilized in activated macrophages (Kobayashi et al., 2016; Mills et al., 2018),, however, our understanding of how Nrf2 regulates macrophage biology in the resting and activated state is incomplete. Our high-resolution proteomics analysis revealed an unexpected role for Nrf2 as a critical regulator of not only redox, but also intermediary metabolism, glycolysis and mitochondrial respiration. This is particularly striking given the importance of metabolic reprogramming for macrophage effector functions (Ryan and O’Neill, 2020). Unexpectedly, we also found that Nrf2 promoted fusion of mitochondrial networks in inflammatory macrophages, which may be due to changes in the abundance of mitochondrial fission/fusion proteins (**Figure S3F**), or alternatively, may be an indirect effect due to altered redox homeostasis, as recently reported in other contexts (Cvetko et al., 2021; Shutt et al., 2012). While it is known that mitochondria signal to Nrf2, these results suggest that activation of Nrf2 has a striking capacity to govern mitochondrial physiology and could have implications for immunoregulatory events.

Excitingly, Nrf2 activation also resulted in a finetuning of the inflammatory response and was found to regulate proteins linked to antigen presentation, T cell activation and cytokine production, which suggests Nrf2 may have an important role in facilitating communication between the innate and adaptive immune systems. Notably, Nrf2 was found to suppress anti-viral response effectors. This finding is of interest due to the paradoxical role of Nrf2 as both an inhibitor of viral replication in certain contexts (Olagnier et al., 2020; Ordonez et al., 2021; Wyler et al., 2019) and a promoter in others (Olagnier et al., 2017). Other cellular stress response pathways, notably the PKR-induced integrated stress response (ISR) and Atf4 lead to a suppression of translation to prevent viral replication (Dauber and Wolff, 2009; Meurs et al., 1990). Given the known interplay of Atf4 and Nrf2 (Kasai et al., 2019), it is possible that Nrf2 may activate similar responses to suppress viral gene expression, even in the presence of impaired anti-viral effectors, and will require further investigation.

Finally, 4-OI has emerged as an anti-inflammatory compound with utility in various disease models via activation of Nrf2 (Li et al., 2020; Liu et al., 2021; Olagnier et al., 2018; Olagnier et al., 2020; Zheng et al., 2020). Importantly, 4-OI represses interferon signaling both in response to viral stimuli and in cases of type I interferonopathies (Olagnier et al., 2018). Here, we confirm a central role for Nrf2 in mediating the immunomodulatory activity of 4-OI in inflammatory macrophages. Considering the interest in Nrf2 activators for treatment of viral infection(s), care must be taken given the divergent response depending on the virus. This emphasizes the importance of future work in elucidating how Nrf2 modulates bacterial and viral infections.

## Methods

### Animals

Mice were bred and maintained at the Medical School Resource Unit of the University of Dundee, with free access to water and food (pelleted RM1 diet from SDS Ltd., Witham, Essex, UK), on a 12-h light/ 12-h dark cycle, 35% humidity. Experimental design was in line with the 3Rs principles of replacement, reduction, and refinement (www.nc3rs.org.uk) and in accordance with the regulations described in the UK Animals (Scientific Procedures) Act 1986 and approved by the Welfare and Ethical use of Animals Committee of the University of Dundee. Wild-type, Nrf2^-/-^: Keap1^+/+^ (Nrf2-KO) and Nrf2^+/+^: Keap1^flox/flox^ (Keap1-KD) mice were on the C57BL/6 genetic background. Both male and female mice were used.

### Generation and treatment of BMDMs

Mice were euthanized in a CO_2_ chamber and death was confirmed by cervical dislocation. Bone marrow cells were extracted from the leg bones and differentiated in DMEM (containing 10% foetal calf serum, 1% penicillin/streptomycin and 20% L929 supernatant) for 6 days, at which time they were counted and re-plated for experiments. Unless stated, 5 × 10^6^ BMDMs per millilitre were used in *in vitro* experiments. The LPS concentration used was 100 ng ml^−1^ (Sigma) and 4-OI was used at 125 μM (synthesized by Dr. Stuart Caldwell and Prof. Richard Hartley).

### Flow cytometry

BMDM cells (2 ×10^5^), isolated and differentiated as described above, were washed with ice-cold DPBS containing 2% FBS (Flow Buffer), re-suspended in 300μl of the Flow Buffer containing 0.2 μg/ml DAPI, acquired on BD LSRFortessa Cell Analyzer and analysed on FlowJo software. Three biologically independent samples were analyzed.

### Western blotting

Following the respective treatments, BMDM cells (5 × 10^5^) grown in 12-well plates were washed twice with PBS before lysing in SDS lysis buffer [50 mM Tris pH 6.8, 10% glycerol (v/v), 2% SDS (w/v) and 0.001% (w/v) Bromophenol Blue]. The samples were sonicated for 30 sec at 20% amplitude before measuring the protein content using the BCA assay (Pierce). Lysates (15 μg total protein) were loaded onto 20-well 4-12% Bis-Tris NuPAGE gel (Thermo) and the proteins were separated by electrophoresis using MOPS buffer (Thermo). Separated proteins were transferred onto 0.45-μm premium nitrocellulose membranes (Amersham) by wet transfer (Biorad). Membranes were blocked in 5% (w/v) non-fat milk (Marvel) dissolved in PBS-0.1% (v/v) Tween-20 (PBST) (Milk-PBST) for 1h. Immunoblotting was performed using primary antibodies generated against GAPDH (1:20000, Proteintech 60004-1-Ig), Keap1 (1:2000, Millipore MABS514), and Nrf2 and Nqo1 (1:1000, Cell Signaling #12721, #62262), all of which were diluted in Milk-PBST. The membranes were incubated with primary antibodies overnight at 4°C or for 1h at RT for GAPDH. Subsequently, the membranes were washed with PBST for 30 min before incubating with either the fluorescently conjugated secondary antibodies (1:20000) (Li-COR) or horseradish-peroxidase conjugated goat anti-rabbit secondary antibody for the Nrf2 blot (1:5000, #7074) (Cell Signaling) for 1 h at RT. Following secondary antibody incubation, the blots were washed for 30 min with PBST before visualizing the proteins by scanning the blots with the Odyssey Clx imager (Li-COR) or detecting the chemiluminescent signal using an X-ray film (Amersham).

### RNA extraction and real-time quantitative (qPCR)

BMDMs (5 × 10^5^ per well) were plated onto 12-well plates, left to adhere overnight and treated as indicated. At experimental endpoint, cells were washed in 1X PBS and then RNA was extracted using RNeasy kit (Qiagen) following the manufacturer’s instructions. RNA was eluted in water and then quantified using a Nanodrop (ThermoFisher Scientific). RNA (1 μg) was reverse-transcribed using High capacity cDNA Reverse Transcription kit (Applied Biosystems). For real-time qPCR, cDNA was run using Fast SYBR^®^ Green Master Mix (Applied Biosystems) according to manufacturer’s instructions and primers were designed for genes of interest (see below) using primer-BLAST. *Rps18* was used as the endogenous control. qPCR experiments were run on a 7900 HT Fast Real-Time PCR System (Applied Biosystems).

### qPCR primers

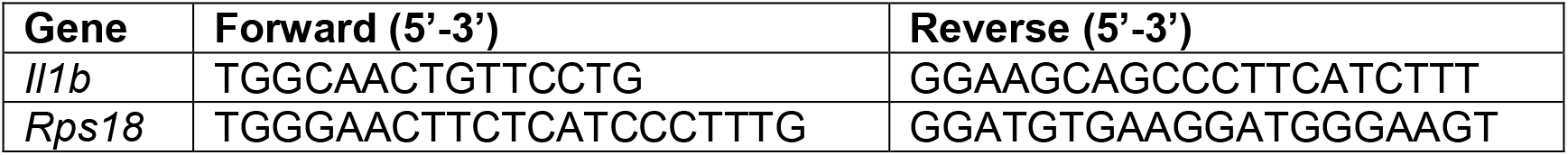

### Proteomic analysis

Sample preparation, tandem mass tag (TMT) labeling, protein digestion, fractionation and peptide LC-MS analysis were performed as described (Howden et al., 2019). Briefly, cell pellets were lysed in 400 μL lysis buffer [4% sodium dodecyl sulfate, 50 mM tetraethylammonium bromide (pH 8.5) and 10 mM tris(2-carboxyethyl)phosphine hydrochloride], lysates were boiled and sonicated before alkylation with 20 mM iodoacetamide for 1 h at 22 °C in the dark. Proteins were digested with LysC and Trypsin, subjected to TMT labeling and the TMT-labelled samples were fractionated using high-pH reverse-phase chromatography. Fractions were dried, peptides (1 μg) dissolved in 5% formic acid and analyzed by LC-MS using an Orbitrap Fusion Tribrid mass spectrometer (Thermo Fisher Scientific) equipped with a Dionex ultra-high-pressure liquid chromatography system (RSLCnano).

### Proteomics data processing

The TMT-labeled samples were collected and analyzed using MaxQuant (Cox and Mann, 2008; Tyanova et al., 2016) v. 1.6.2.10. The TMT channel mapping is shown in Supplementary File 1. The FDR threshold was set to 1% for each of the respective Peptide Spectrum Match (PSM) and Protein levels. The data were searched with the following parameters; type was set to Reporter ion MS3 with 10 plex TMT, stable modification of carbamidomethyl (C), variable modifications, oxidation (M), acetylation (protein N terminus), deamidation (NQ), with a 2 missed tryptic cleavages threshold. Minimum peptide length was set to 6 amino acids. Proteins and peptides were identified using Uniprot (SwissProt May 2018). Run parameters have been deposited to PRIDE (Perez-Riverol et al., 2019) along with the full MaxQuant quantification output (PXD027737). All corrected TMT reporter intensities were normalized and quantified to obtain protein copy number using the proteomic ruler method (Wisniewski et al., 2014) as described in (Howden et al., 2019).

### LC-MS Metabolomics

#### Steady-state metabolomics

For steady-state metabolomics, 5 × 10^5^ cells were plated the day before onto 12-well plates (5 technical replicates from 3 biological replicates) and extracted at the appropriate experimental endpoint (24 h timepoint). Prior to metabolite extraction, cells were counted using a hemocytometer using a separate counting plate prepared in parallel and treated exactly like the experimental plate. At the experimental endpoint, the media was aspirated off and the cells were washed at room temperature with PBS and placed on a cold bath with dry ice. Metabolite extraction buffer (MES) was added to each well following the proportion 1×10^6^ cells/0.5 ml of buffer. After 10 minutes, the plates were stored at −80°C freezer and kept overnight. The following day, the extracts were scraped and mixed at 4°C for 15 min in a thermomixer at 2000 rpm. After final centrifugation at max speed for 20 min at 4°C, the supernatants were transferred into labelled LC-MS vials.

#### Liquid chromatography coupled to Mass Spectrometry (LC-MS) analysis

HILIC chromatographic separation of metabolites was achieved using a Millipore Sequant ZIC-pHILIC analytical column (5 μm, 2.1 × 150 mm) equipped with a 2.1 × 20 mm guard column (both 5 mm particle size) with a binary solvent system. Solvent A was 20 mM ammonium carbonate, 0.05% ammonium hydroxide; Solvent B was acetonitrile. The column oven and autosampler tray were held at 40°C and 4°C, respectively. The chromatographic gradient was run at a flow rate of 0.200 mL/min as follows: 0–2 min: 80% B; 2-17 min: linear gradient from 80% B to 20% B; 17-17.1 min: linear gradient from 20% B to 80% B; 17.1-22.5 min: hold at 80% B. Samples were randomized and analyzed with LC–MS in a blinded manner with an injection volume was 5 μl. Pooled samples were generated from an equal mixture of all individual samples and analyzed interspersed at regular intervals within sample sequence as a quality control.

Metabolites were measured with a Thermo Scientific Q Exactive Hybrid Quadrupole-Orbitrap Mass spectrometer (HRMS) coupled to a Dionex Ultimate 3000 UHPLC. The mass spectrometer was operated in full-scan, polarity-switching mode, with the spray voltage set to +4.5 kV/-3.5 kV, the heated capillary held at 320°C, and the auxiliary gas heater held at 280 °C. The sheath gas flow was set to 25 units, the auxiliary gas flow was set to 15 units, and the sweep gas flow was set to 0 unit. HRMS data acquisition was performed in a range of *m/z* = 70–900, with the resolution set at 70,000, the AGC target at 1 × 10^6^, and the maximum injection time (Max IT) at 120 ms. Metabolite identities were confirmed using two parameters: (1) precursor ion m/z was matched within 5 ppm of theoretical mass predicted by the chemical formula; (2) the retention time of metabolites was within 5% of the retention time of a purified standard run with the same chromatographic method. The acquired spectra were analysed using XCalibur Qual Browser and XCalibur Quan Browser software (Thermo Scientific) and the peak area for each detected metabolite was normalized against the total ion count (TIC) of that sample to correct any variations introduced from sample handling through instrument analysis. The normalized areas were used as variables for further statistical data analysis.

### Oxygen consumption rate (OCR) and extracellular acidification rate (ECAR) measurements

Oxygen consumption rate (OCR) and extracellular acidification rate (ECAR) were measured using the real-time flux analyzer Seahorse XF24 (Agilent) according to a method modified from Van den Bossche et al. (Van den Bossche et al., 2015). In brief, 0.5 × 10^5^ cells were plated onto the instrument cell plate 27 h before the experiment in complete RPMI 1640 medium (Invitrogen) supplemented with 10% FBS, 2mM L-glutamine and 1mM Na-pyruvate (5 replicate wells for each condition). Following adhesion, the cells were treated as indicated for 24 h. At the treatment endpoint, the cell culture medium was replaced with XF RPMI medium pH 7.4 (Agilent, 103576-100) supplemented with 2 mM glutamine prior to analysis. Cells were treated with 25 mM glucose, 1.5 μM oligomycin, 1.5 μM FCCP/1mM Na-pyruvate and 2.5 μM antimycin A/1.25 μM rotenone to assess the respiration parameters.

### Confocal microscopy

#### Mitochondrial morphology

BMDMs were fixed with 4% (w/v) PFA in PBS for 15 min, 37°C, 5% CO2 and then washed three times with PBS. Autofluorescence was quenched with 50 mM NH_4_Cl for 10 min at room temperature, followed by three washes with PBS. BMDMs were permeabilized with 0.1% (v/v) TritonX-100 in PBS for 10 min at room temperature. The permeabilized cells were then blocked with 10% FBS in PBS for 20 min at room temperature. BMDMs were incubated in rabbit anti- toM20 antibody (Proteintech, 11802-1-AP) at 1/1000 dilution in 5% FBS in PBS for 2 hours at room temperature followed by three washes in 5% FBS in PBS. Cells were then incubated in goat anti-rabbit Alexa 569 antibody (Invitrogen, A11036) at 1/1000 dilution in 5% FBS for 1 h at room temperature. BMDMs were washed three times with PBS and then stored in PBS at 4°C until imaging.

BMDMs were imaged using a 100x oil objective lens with 500 ms exposure time, 50% laser intensity using excitation/emission wavelengths 561/620-60 nm on an Andor Spinning Disk confocal microscope. Images were analysed using Fiji ImageJ. Mitochondrial morphology was assigned as intermediate, fused/elongated or fragmented and presented as mean % of all cells ± SEM. 75 cells were counted for each condition, for 3 mice.

Mitochondrial morphological parameters were measured as number of mitochondria, average length of mitochondria and average area of mitochondria in 20 regions of interest (ROIs) for each condition, for 3 mice. The ROI is a 15×15 um box that is chosen near the edge of cells where mitochondrial networks are more clearly defined, but chosen from random cells.

### Statistical analysis

For metabolomics data, metaboanalyst 5.0 (Pang et al., 2021) was used to analyze, perform statistics and visualize the results. One-way ANOVA corrected for multiple comparisons by the Tukey statistical test was used and a p. Adjusted < 0.05 was set as the cut-off. For proteomics data, protein copy number was converted to a log_2_ scale and biological replicates were grouped by experimental condition. Protein-wise linear models combined with empirical Bayes statistics were used for the differential expression analyses. The Bioconductor package limma was used to carry out the analysis using an R based online tool (Shah et al., 2020). Data were visualized using a volcano plot, which shows the log_2_ fold change on the x axis and the adjusted p value on the y axis. Overrepresentation analysis (ORA) of significant changes were assessed using Enrichr (Kuleshov et al., 2016) and the Bioconductor package clusterProfiler 4.0 in R (version 3.6.1). Emapplots were generated using enrichplot package in R (version 3.6.1). Graphpad Prism 8.0 was used to calculate statistics in bar plots.

## Contributions

D.G.R – designed and performed experiments for proteomics and metabolomics, provided intellectual input, analyzed and visualized the data, prepared the figures and co-wrote the paper. E.V.K – harvested tissues, designed and performed experiments for respirometry analysis and flow cytometry, provided intellectual input, analyzed and visualized data, and participated in manuscript writing. A.C – performed analysis of mitochondrial morphology using confocal microscopy. J.L.H. and A.J.B. performed the proteomics analysis. S.D.N and M.H. performed immunoblotting analysis, animal breeding and genotyping. L.T. and E.N. assisted with metabolomics. L.A.J.O., C.F., A.J.L., A.Y.A., D.A.C., and M.P.M. provided expert intellectual input and oversaw various aspects of the work. A.T.D.K - conceptualized the project, oversaw the work, and co-wrote the manuscript. All authors appraised and edited the manuscript.

## Acknowledgments

We thank Cancer Research UK (C20953/A18644) and the Wellcome Trust (105024/Z/14/Z) for funding. This work was also supported by a UK Research Partnership Infrastructure Fund award to the Centre for Translational and Interdisciplinary Research. We would like to thank Dr. Lisa Dwane and Dr. Christina Schmidt for discussions around and assistance with data visualization, and Dr Stuart Caldwell and Prof. Richard Hartley who originally synthesized 4-octyl itaconate (4-OI), see (Mills et al., 2018). We would also like to thank Dr. Vincent Paupe and Dr. Roy Chowdhury for advice with mitochondrial imaging and terminology.

---

**Figure S1.**
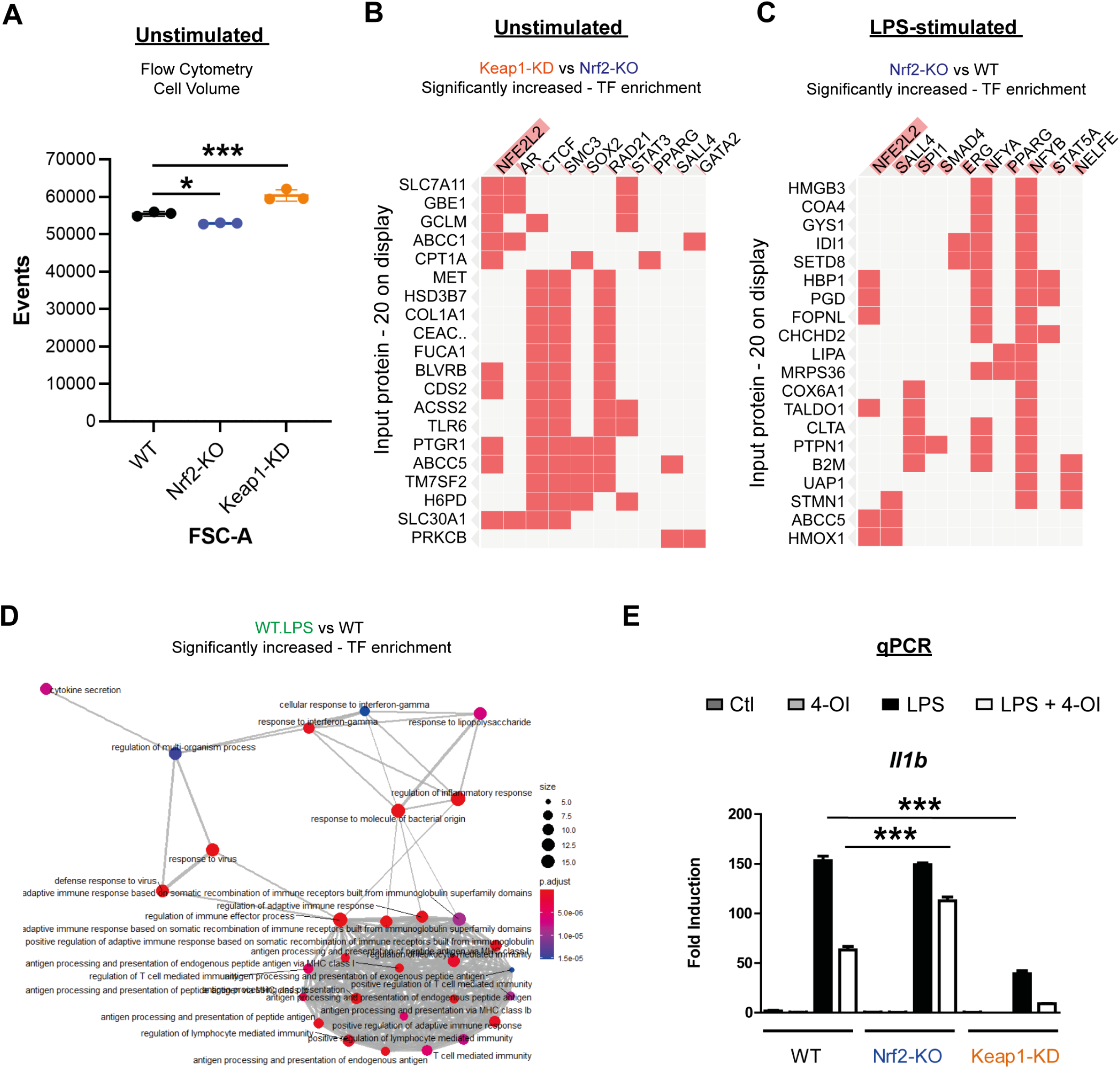
Supporting data for figure 1. **(A)** Flow cytometry analysis of cell size in unstimulated state **(B)** TF enrichment of decreased targets in unstimulated Keap1-KD compared to Nrf2-KO **(C)** TF enrichment of decreased targets in LPS stimulated Nrf2-KO compared to WT **(D)** Enrichment map of GO: biological processes of significantly increased proteins with LPS stimulation in WT **(E)** qPCR (n = 3 biological replicates) validation of 4-OI (125 μM) inhibition of IL-1β **(A, E)** (n = 3 biological replicates). Data are mean ± SEM. p value determined by one-way ANOVA, corrected for multiple comparisons by Tukey statistical test. p <0.05*; p <0.01**; p < 0.001***. **(D)** ORA by clusterProfiler, FDR correction by Bonferroni test **(B-C)** Determined by Enrichr using ENCODE and ChEA database (combined score – p value and z score).

**Figure S2.**
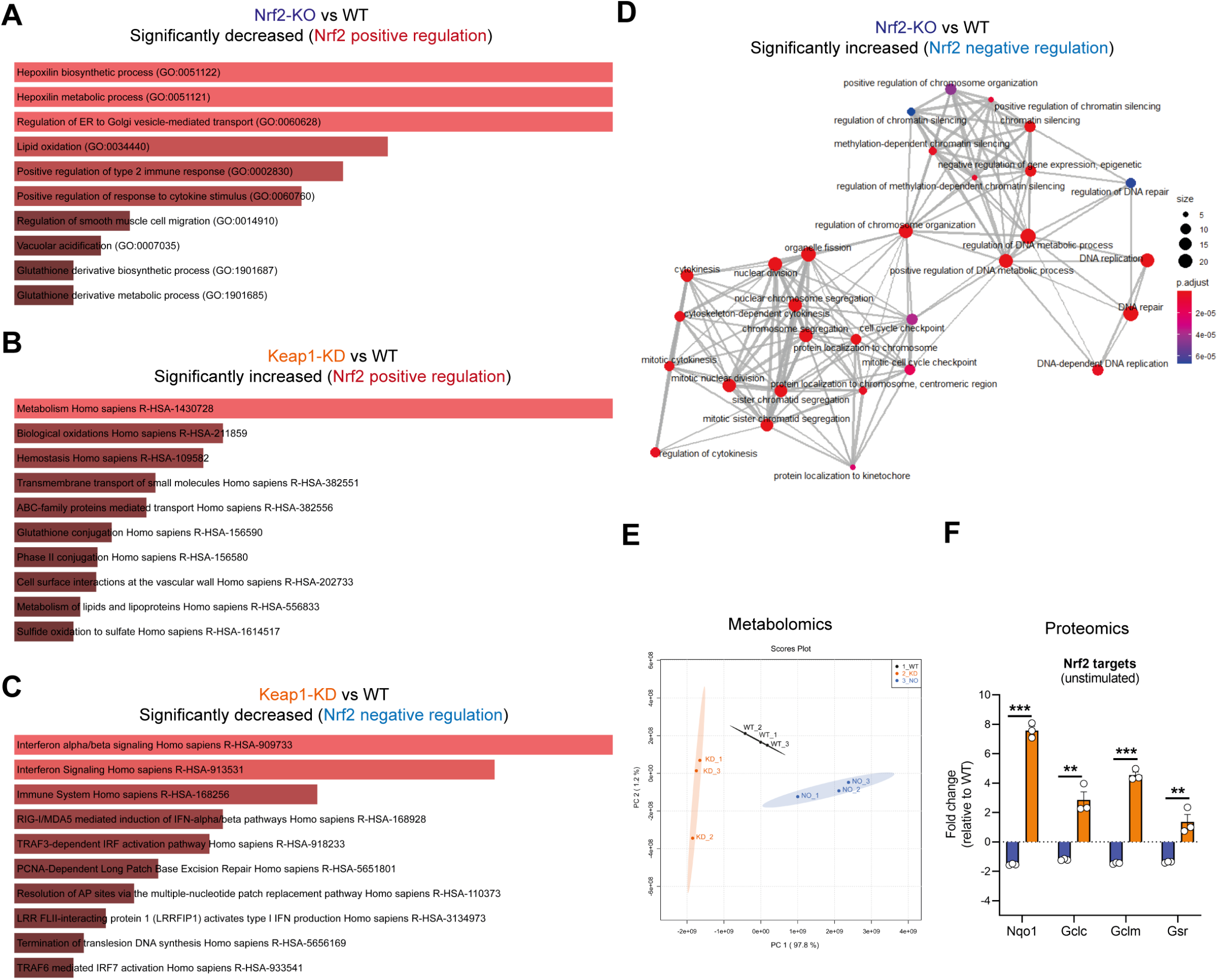
Supporting data for figure 2. **(A)** Enrichment of Reactome pathways of Nrf2 positively regulated targets (Nrf2-KO vs WT) **(B)** Enrichment of Reactome pathways of Nrf2 positively regulated targets (Keap1-KD vs WT) **(C)** Enrichment of Reactome pathways of Nrf2 negatively regulated targets (Keap1-KD vs WT) **(A-C)** ORA by Enrichr and FDR correction by Bonferroni test. **(D)** Enrichment map of GO: biological processes of Nrf2 negatively regulated targets (Nrf2-KO vs WT). ORA by clusterProfiler, FDR correction by Bonferroni test. **(E)** PCA plot of metabolomics. Determined by Metaboanalyst 5.0. **(F)** Fold change of prototypical Nrf2 targets (Nrf2-KO vs WT or Keap1-KD vs WT). Data are mean ± SEM. p value determined by multiple t tests - one per row, corrected for multiple comparisons by Holm-Sidak test. p <0.05*; p <0.01**; p < 0.001***.

**Figure S3.**
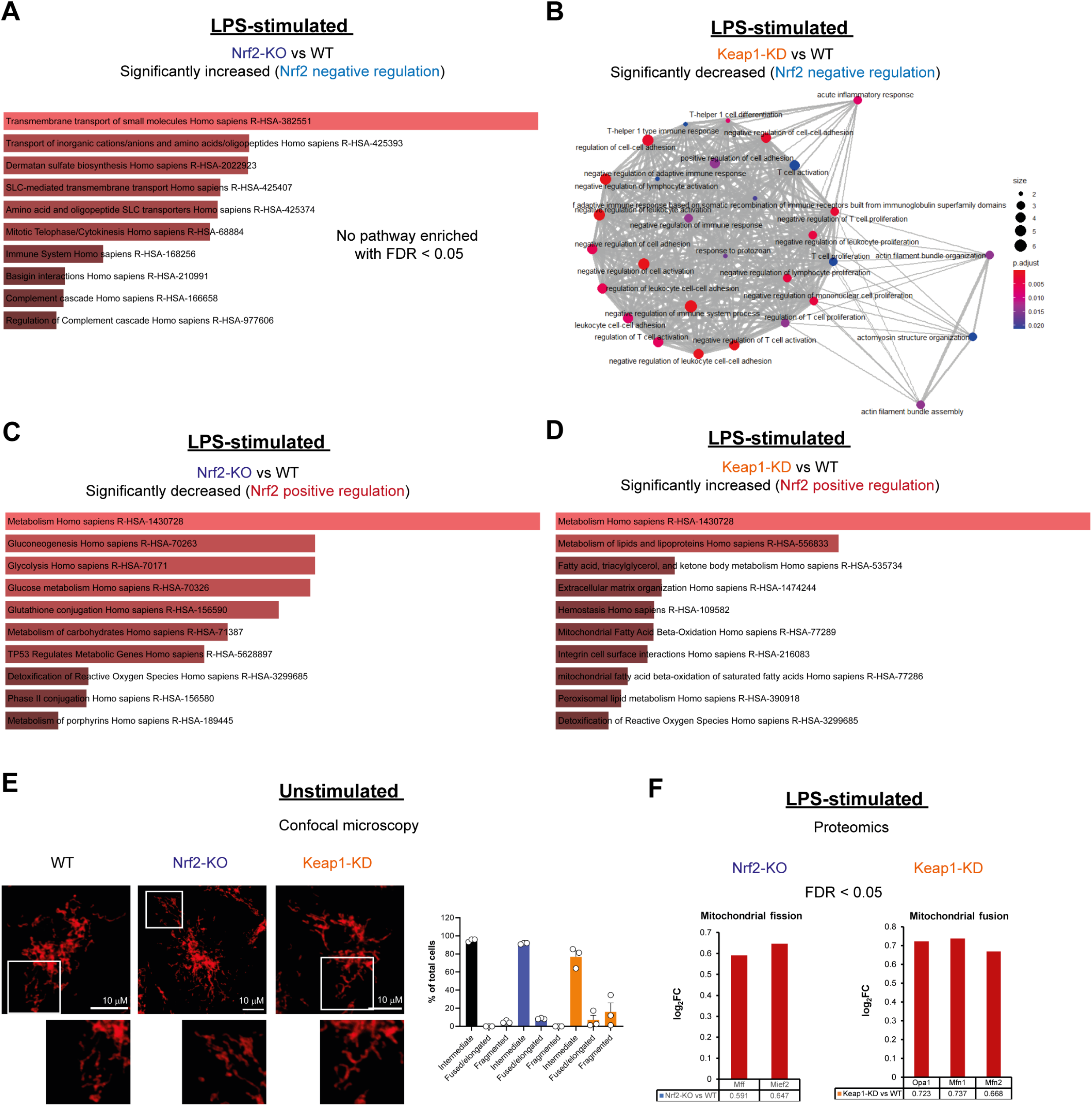
Supporting data for figure 3. **(A)** Enrichment map of GO: biological processes of Nrf2 negatively regulated targets (Nrf2-KO vs WT) **(B)** Enrichment map of GO: biological processes of Nrf2 negatively regulated targets (Keap1-KD vs WT) **(A-B)** ORA by clusterProfiler, FDR correction by Bonferroni test. **(C)** Enrichment of Reactome pathways of Nrf2 positively regulated targets (Nrf2-KO vs WT) **(D)** Enrichment of Reactome pathways of Nrf2 positively regulated targets (Keap1-KD vs WT) **(C-D)** ORA by Enrichr and FDR correction by Bonferroni test. **(E)** Confocal microscopy of mitochondrial morphology using Tom20 (images are representative, bar plot n = 3 biological replicates). Data are mean ± SEM. p value determined by one-way ANOVA, corrected for multiple comparisons by Tukey statistical test. **(F)** Log_2_FC from LPS stimulated proteomics (Nrf2 vs WT & Keap1-KD vs WT).

**Figure S4.**
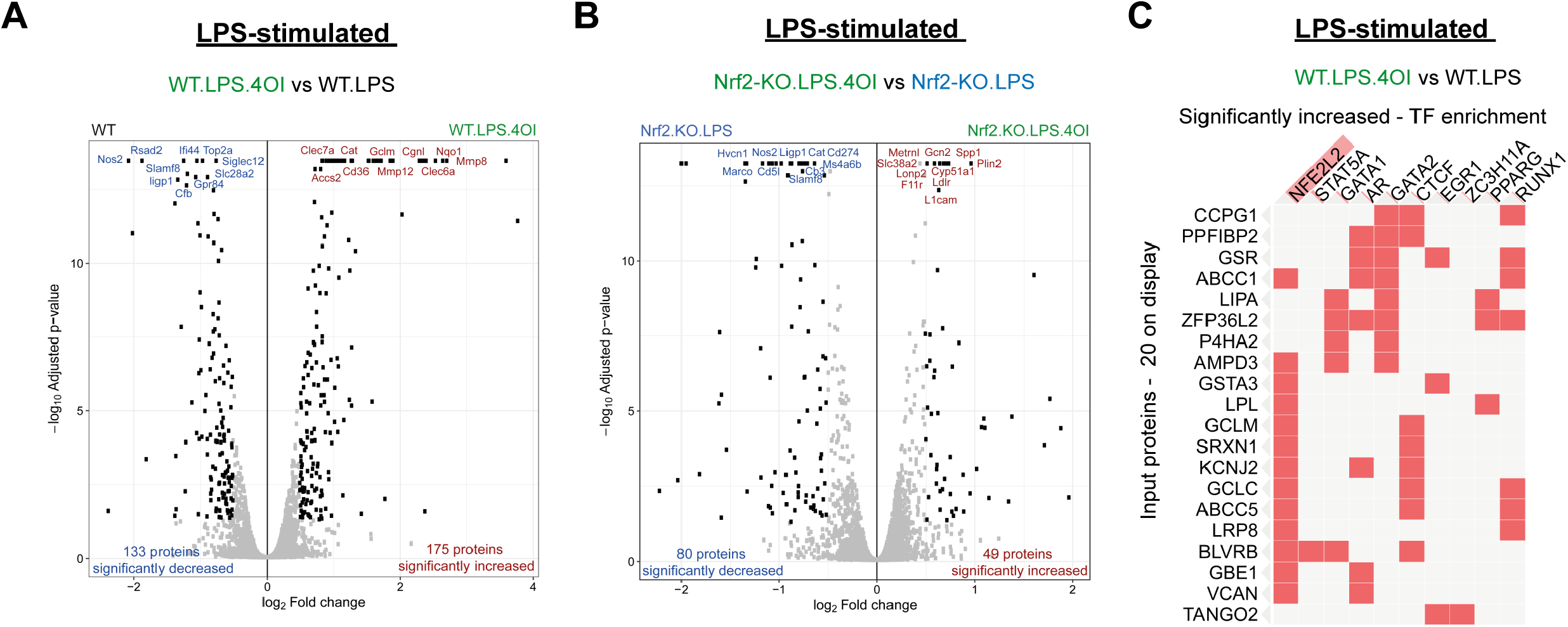
Supporting data for figure 4. **(A)** Volcano plot of LPS with 4-OI compared to LPS only (WT) **(B)** Volcano plot of LPS with 4-OI compared to LPS only (Nrf2-KO) **(C)** TF enrichment of increased targets in LPS with 4-OI compared to LPS only (WT). **(A-B)** ORA by Enrichr, FDR correction by Bonferroni test. **(C)** Determined by Enrichr using ENCODE and ChEA database (combined score – p value and z score).

## Notes

### Competing Interest Statement

The authors have declared no competing interest.

